# Barcode-integrated reverse transcription for accurate, complete, and low-input RNA sequencing

**DOI:** 10.64898/2026.07.15.738812

**Authors:** Wei Chen, Yizhou Zhu, Xiaoyu Weng, Qi Jiang, Qiong Zhao, Joshua Than, Moshe Baruch, David Yu Zhang

## Abstract

In conventional total-RNA sequencing, unwanted RNA is removed by ribosomal-RNA depletion or messenger-RNA enrichment, and contaminating genomic DNA is cleared by DNase, both before reverse transcription. A sample index is added only later, during amplification, so each sample is carried through reverse transcription and cleanup on its own. These steps lose material that low-input samples cannot spare, and the per-sample handling limits throughput. Here we describe BIRT (Barcode-Integrated Reverse Transcription), a hairpin primer that carries a sample barcode and begins reverse transcription preferentially at RNA 3′ ends, folding both barcoding and 3′-end capture into first-strand synthesis. Barcoding at reverse transcription enables early pooling, so samples are combined before any cleanup. It also authenticates RNA against DNA contamination: the primer barcodes RNA 909-fold more often than genomic DNA, and non-barcoded reads, which report DNA, are discarded. The 3′-end preference recovers terminal sequence that random priming loses: for a typical small nucleolar RNA, 66% of reads begin within two nucleotides of the mature 3′ end, against a few percent for random hexamers. We pair BIRT with PERD (Probes for Excess RNA Depletion), a modular set of blocking probes that removes unwanted RNA within the same reaction without distorting expression (Pearson r = 0.98). We have applied BIRT to 8,079 samples across seven species, from cultured cells and primary tissue to FFPE and extracellular-vesicle RNA.

## Introduction

Bulk RNA sequencing underlies most of transcriptomics [1], yet several of its recurring limitations trace to a single fact about library construction: some information is lost at reverse transcription, and the rest of the preparation is handled in separate steps around it. Two kinds of information are lost at reverse transcription itself. It is primed with oligo(dT), which captures only polyadenylated RNA [7], or with random hexamers, which start at internal positions and discard the 3′-terminal sequence [2], so the position of a transcript’s end goes unrecorded. Separately, RNA and any co-purified genomic DNA are copied into the same unmarked cDNA and become indistinguishable; DNase treatment before reverse transcription is the usual defense but, as a separate step, is rarely complete for precious or degraded samples [14]. The rest of the preparation is handled around reverse transcription. The sample index is added late, during or after amplification, so each sample is reverse-transcribed and cleaned separately. Every cleanup loses a fixed fraction of cDNA, which low-input material can least afford. Ribosomal RNA, which dominates total RNA, is removed by a further depletion step before reverse transcription [6], adding another cleanup with its own loss and bias.

These are usually treated as four independent problems with four independent fixes. We reasoned instead that recording the missing information in the cDNA at the moment of reverse transcription could address them together. Here we describe BIRT (Barcode-Integrated Reverse Transcription), a hairpin reverse-transcription primer that carries a sample barcode and initiates preferentially at RNA 3′ ends, and PERD (Probes for Excess RNA Depletion), which removes unwanted RNA within the same reaction. Barcoding during first-strand synthesis, an approach used to mark molecular origin in single-cell transcriptomics [3–5], lets samples be pooled before any cleanup and marks every RNA-derived read, so that contaminating DNA is removed after sequencing; with PERD, ribosomal and other unwanted RNA are depleted in the same step. We show that BIRT and PERD anchor reverse transcription to transcript 3′ ends, separate RNA from DNA contamination, raise throughput and low-input sensitivity through early pooling, deplete unwanted RNA without distorting expression, and operate across species and sample types at scale.

## Results

### 3′-end anchoring of reverse transcription

Random-hexamer priming initiates reverse transcription at internal positions, so the 3′-terminal sequence of a transcript is frequently absent from the resulting cDNA (Fig. 1a). To direct priming to the transcript end, we replaced the free hexamer with a hairpin primer that carries a random hexamer at its 5′ end, a universal double-stranded stem, and a sample barcode in the loop (Fig. 1b). We hypothesized that when the hexamer anneals at a 3′ terminus, the RNA:hexamer duplex stacks coaxially with the universal stem and stabilizes the primed complex, whereas at an internal site the downstream single-stranded RNA sterically clashes with and electrostatically repels the hairpin and cannot stack, so terminal priming is favored (Fig. 1b).

**Figure 1.**
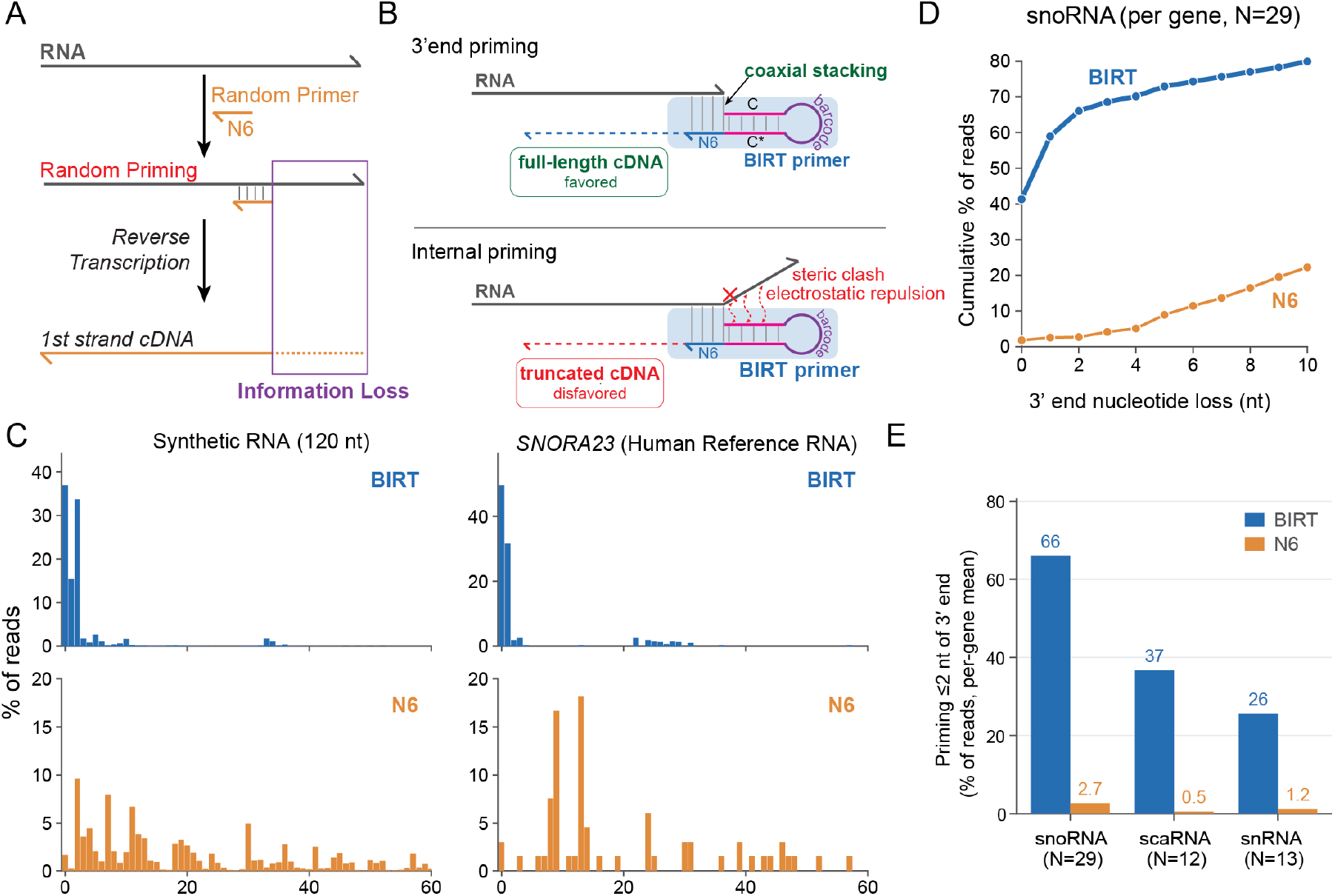
BIRT anchors reverse transcription to RNA 3′ ends. (a) Random-hexamer priming initiates reverse transcription at internal positions, losing the 3′-terminal sequence of the transcript. (b) The BIRT primer is a hairpin carrying a random hexamer at its 5′ end, a universal stem, and a sample barcode in the loop; at a transcript 3′ end the RNA:hexamer duplex stacks coaxially with the stem and terminal priming is favored, whereas at an internal site the downstream RNA forms a single-stranded flap that sterically clashes with and electrostatically repels the hairpin and cannot stack. (c) Distance of the priming end from the 3′ terminus for BIRT (top) and random hexamer (N6, bottom), on a synthetic 120-nt RNA and on the human small nucleolar RNA SNORA23. (d) Per-gene cumulative distribution of the distance to the annotated 3′ end for snoRNAs (N = 29 genes), BIRT versus N6. (e) Fraction of reads priming within 2 nt of the 3′ end (per-gene mean) for snoRNA (N = 29), scaRNA (N = 12), and snRNA (N = 13), BIRT versus N6. The snRNA value is a lower bound because reference 3′ annotations extend beyond the mature terminus.

To test this prediction, we first measured priming positions on defined templates (Fig. 1c). On a synthetic 120-nt RNA, the BIRT primer placed 86% of priming events within 2 nt of the 3′ terminus, against 11% for a free random hexamer; on the endogenous small nucleolar RNA SNORA23 in cellular RNA, the corresponding fractions were 83% and 3%. We then extended the measurement transcriptome-wide. Because a single hyper-abundant snoRNA (U3) dominates snoRNA reads and anchors less tightly than the rest, we computed the fraction of reads within 2 nt of the annotated 3′ end per gene and averaged across genes (Fig. 1d). By this per-gene measure, the typical snoRNA reached 66% within 2 nt with BIRT against 2.7% with random priming, and small Cajal body RNAs reached 37% against 0.5% (Fig. 1e). Small nuclear RNAs reached 26% against 1.2%; this is likely an underestimate, because their reference 3′ annotations extend beyond the mature terminus. Messenger RNAs and lncRNAs showed no enrichment at the annotated 3′ end. Because these transcripts are long and are randomly fragmented before reverse transcription, the mature terminus is only a small fraction of their molecules and each fragment carries its own 3′ end; BIRT still primes preferentially at these fragment ends, but the ends are set by random fragmentation rather than by the transcript terminus, so priming is distributed along the transcript body. BIRT therefore captures the mature 3′ end of transcripts that have a discrete one, while giving full-length body coverage of long, fragmented transcripts.

### Separation of RNA signal from DNA contamination

A total-RNA library cannot distinguish a read derived from RNA from one derived from co-purified genomic DNA, because a random primer copies both into the same unmarked cDNA (Fig. 2a). The BIRT primer addresses this in two ways. First, it is intrinsically RNA-selective: across matched input it barcoded 89.1% of RNA-derived molecules but only 0.098% of genomic DNA, a 909-fold preference (Fig. 2b). Second, because only RNA-templated cDNA receives a barcode, DNA-derived reads can be removed after sequencing by retaining only reads that carry a valid BIRT barcode, a step we refer to as barcode gating (Fig. 2d).

**Figure 2.**
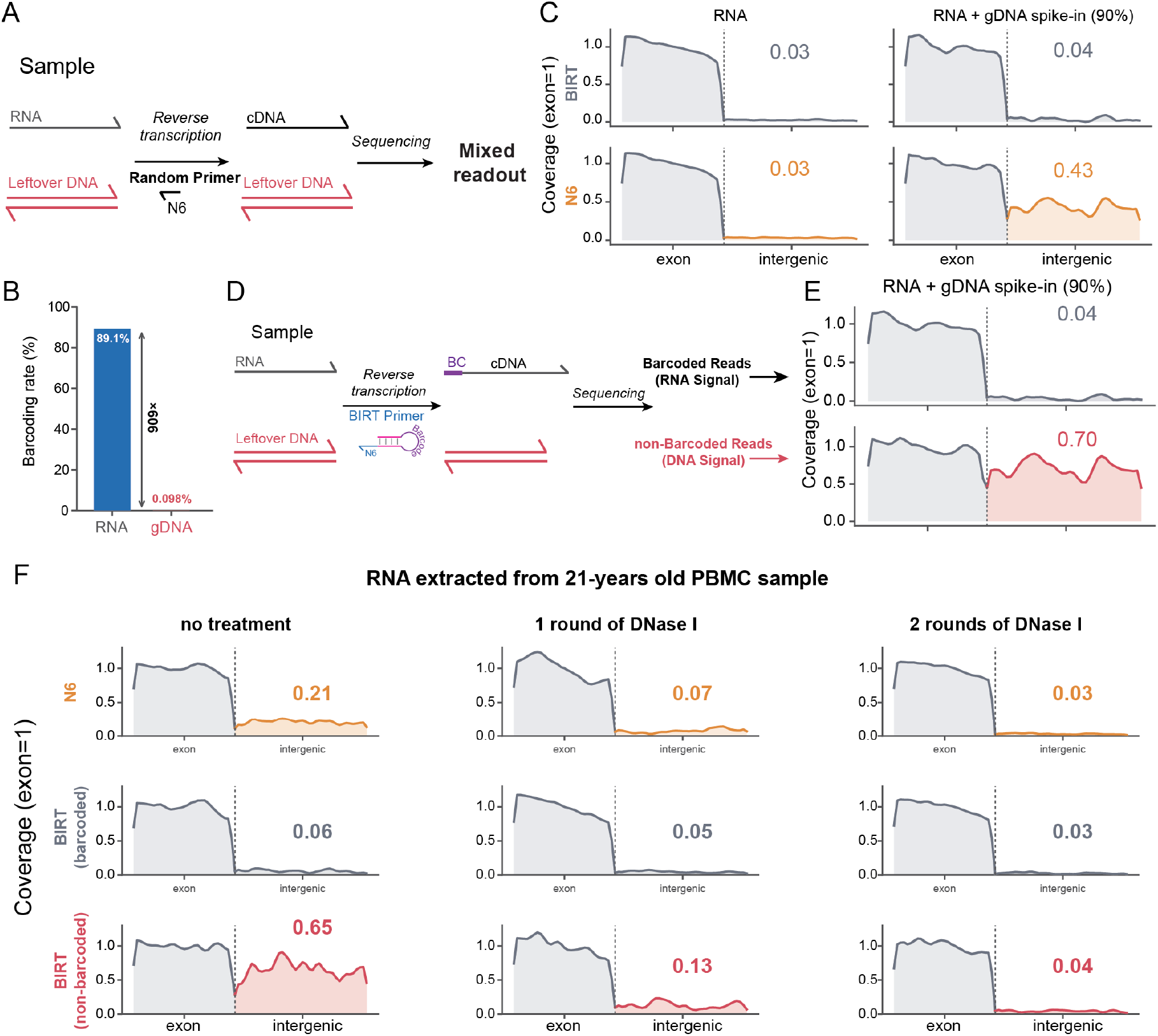
A barcode written at reverse transcription separates RNA signal from DNA contamination. (a) A random primer copies both RNA and co-purified genomic DNA into unmarked cDNA, giving a mixed readout. (b) Barcoding rate of the BIRT primer on RNA versus genomic DNA (909-fold higher for RNA). (c) Coverage over intergenic regions relative to exon for BIRT (top) and N6 (bottom), in RNA and in RNA with a 90% genomic-DNA spike-in. (d) Only RNA-templated cDNA carries a barcode, so barcoded reads report RNA signal and non-barcoded reads report leftover DNA. (e) Intergenic coverage of the barcoded and non-barcoded BIRT fractions in the genomic-DNA spike-in. (f) Intergenic coverage for N6, barcoded BIRT, and non-barcoded BIRT in RNA extracted from a 21-year-old PBMC sample, without treatment and after one or two rounds of DNase I.

To test the first property, we spiked genomic DNA into reference RNA, prepared libraries with BIRT or a random hexamer, and measured coverage over intergenic regions relative to exon (Fig. 2c). In pure RNA both methods were clean (intergenic coverage ≈ 0.03). With a 90% genomic-DNA spike, the random-primed library acquired strong intergenic coverage (0.43), whereas the BIRT library remained at pure-RNA levels (0.04). To test the second property, we separated the BIRT reads by barcode: the barcoded fraction stayed clean (0.04) while the non-barcoded fraction rose to 0.70, tracking the contaminating DNA (Fig. 2e). We then applied barcode gating to a demanding real sample, RNA extracted from a 21-year-old PBMC specimen, with zero, one, or two rounds of DNase I treatment (Fig. 2f). Without any DNase, the random-primed library and the non-barcoded BIRT fraction carried substantial intergenic coverage (0.21 and 0.65), whereas the barcoded BIRT fraction was already clean (0.06) and reached the same floor as two rounds of DNase (0.03 to 0.04). Barcode gating therefore acts as a computational DNase, authenticating RNA-derived reads after sequencing rather than requiring complete enzymatic removal before it.

### Early pooling and low-input detection

In conventional preparation each sample is reverse-transcribed and cleaned on its own, so the fixed fraction of cDNA lost to plastic during cleanup is paid once per sample; for low-input samples this loss removes a large share of the material and, with it, real transcripts (Fig. 3a). Because BIRT attaches the barcode during first-strand synthesis, samples can be pooled immediately after reverse transcription and the whole pool cleaned once, incurring the fixed loss a single time.

**Figure 3.**
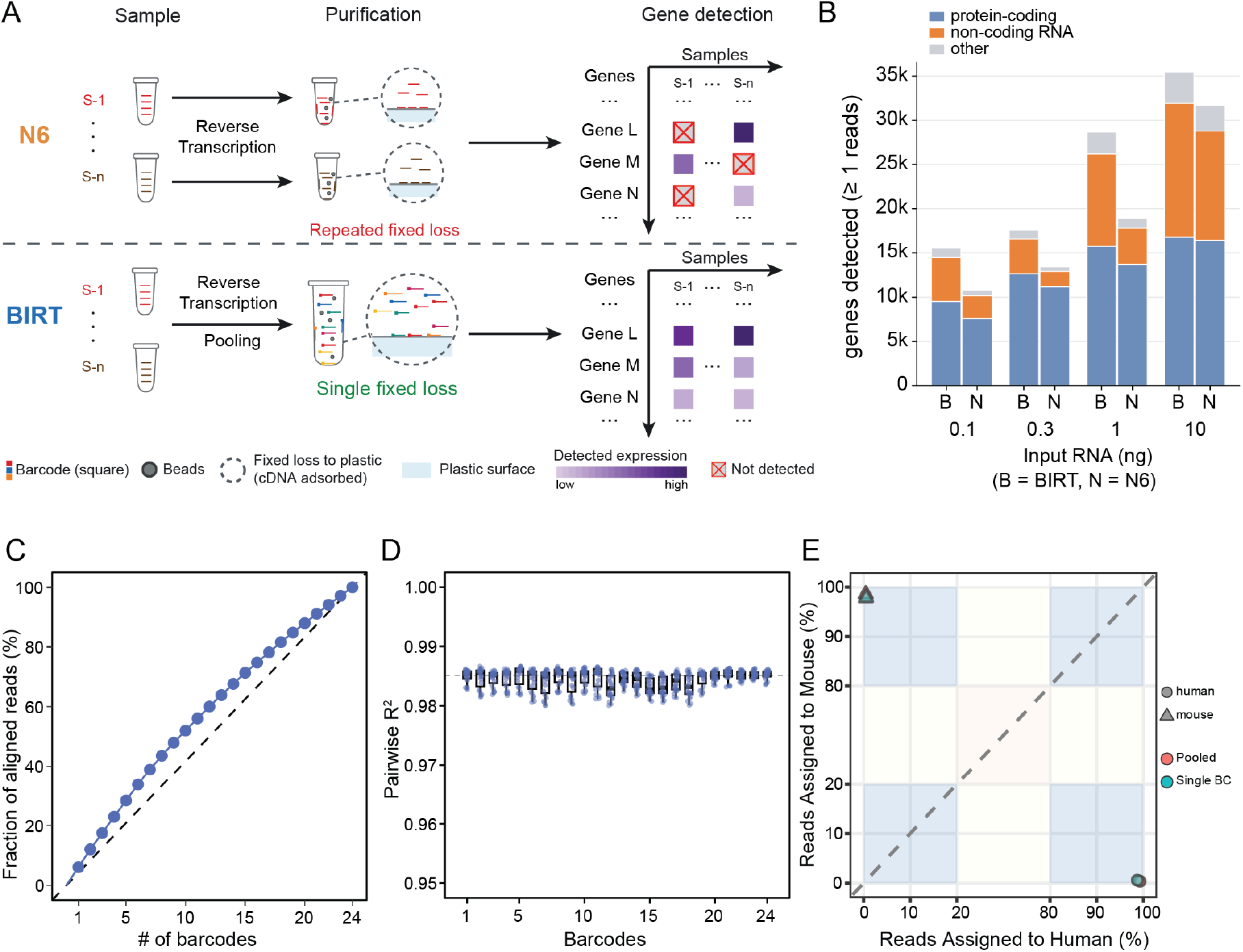
Early pooling raises low-input detection without barcode bias. (a) Conventional preparation reverse-transcribes and cleans each sample separately, repeating the fixed cDNA loss once per sample and detecting fewer low-expression genes; BIRT barcodes during first-strand synthesis and pools before a single cleanup, incurring the loss once. (b) Genes detected (≥ 1 read) versus input RNA (0.1 to 10 ng) by biotype (protein-coding, non-coding RNA, other), for BIRT (B) and random hexamer (N). (c) Fraction of aligned reads versus number of pooled barcodes (1 to 24). (d) Pairwise cross-barcode reproducibility of expression across 24 barcodes. (e) Barnyard species assignment for pooled and single-barcode human/mouse libraries.

To measure the effect on detection, we titrated input RNA from 10 ng to 0.1 ng and compared BIRT with a random-hexamer control at matched depth (Fig. 3b). BIRT detected more genes than the control at every input, and the advantage widened as input fell and was largest for the non-coding fraction, the low-abundance transcripts most vulnerable to per-sample loss. We next asked whether pooling or the barcode itself distorts the data. Pooling one to twenty-four barcodes, the fraction of aligned reads grew smoothly and sub-linearly with pool size (Fig. 3c). To test whether the barcode sequence biases quantification, we pooled twenty-four replicate samples carrying different barcodes and found expression highly reproducible across barcodes (pairwise R^2^ ≈ 0.98; Fig. 3d). To test for cross-sample leakage between pooled barcodes, we constructed a mixed human and mouse library; demultiplexed reads aligned to their species of origin with greater than 99% specificity, indicating negligible barcode crosstalk (Fig. 3e). Early pooling therefore raises detection, most for the hardest-to-detect transcripts, without introducing barcode-dependent bias.

### Depletion of unwanted RNA by PERD

Some RNA species dominate a library without carrying information of interest, most prominently ribosomal RNA, and are typically removed to raise the effective sequencing yield; but enzymatic and probe-based depletion are separate steps performed before reverse transcription, each adding a cleanup with its own loss and potential bias [6]. To fold depletion into the same reaction, we designed PERD (Probes for Excess RNA Depletion), a modular set of blocking probes that hybridize to a target RNA and prevent the barcoded primer from extending along it, while on the RNA of interest the primer displaces the probes and proceeds normally (Fig. 4a). Because PERD requires no enzyme, only a few minutes of added hybridization, and no separate cleanup, it preserves throughput relative to RNase-H-based depletion (Fig. 4b).

**Figure 4.**
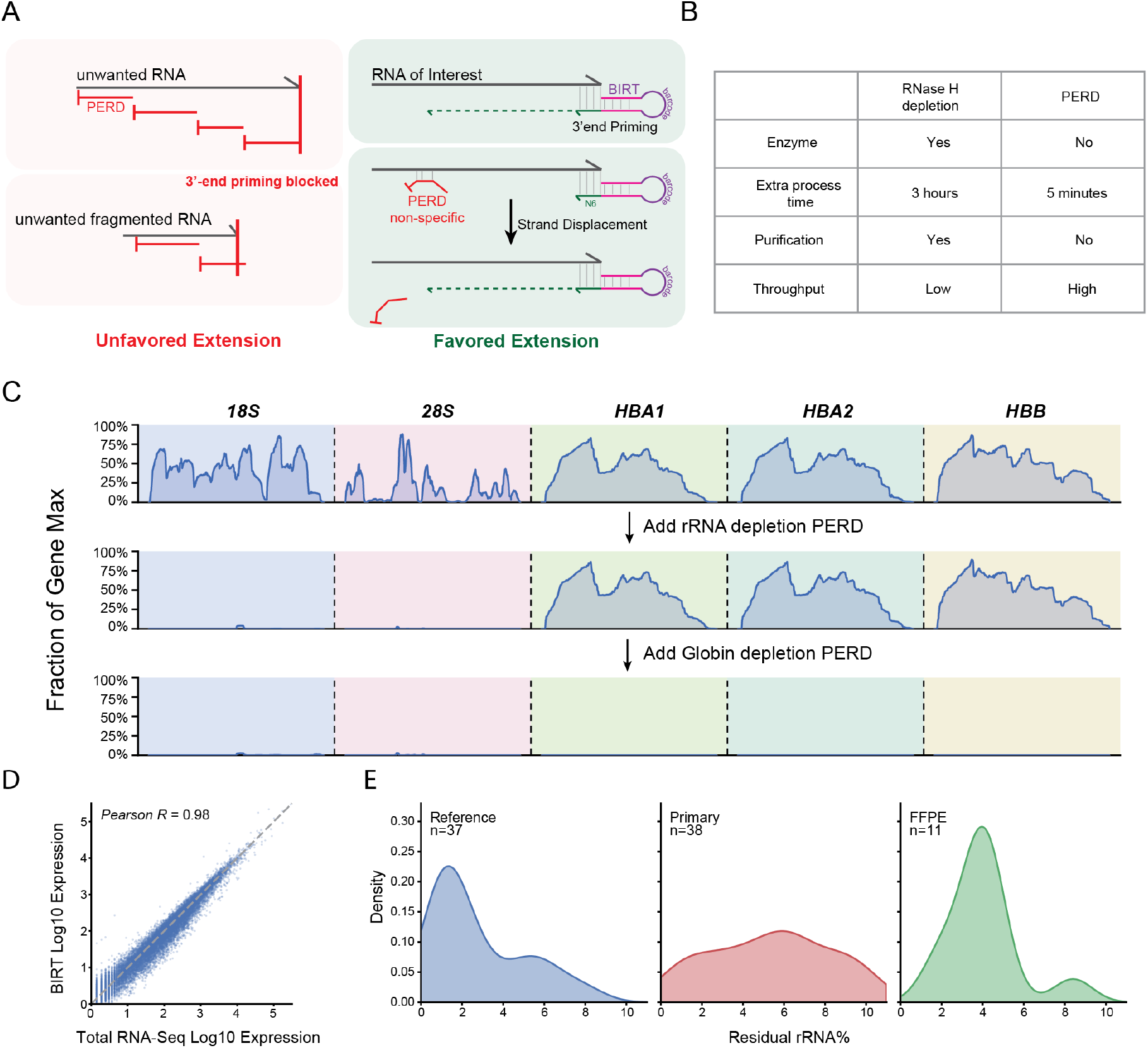
PERD depletes unwanted RNA within the reverse-transcription step. (a) PERD blocking probes hybridize to unwanted RNA and block 3′-end priming and extension (unfavored), whereas on the RNA of interest the BIRT primer primes and displaces the probe by strand displacement (favored). (b) PERD versus RNase-H-based depletion: PERD requires no enzyme, minimal added time, and no separate purification, giving higher throughput. (c) Coverage (fraction of each gene’s maximum) over 18S and 28S rRNA and over globin genes (HBA1, HBA2, HBB), without depletion (top), after adding the rRNA depletion probes (middle; rRNA depleted, globin retained), and after further adding globin depletion probes (bottom; both depleted). (d) Per-gene expression with BIRT versus total RNA-seq (Pearson r = 0.980). (e) Residual ribosomal-RNA fraction for reference (n = 37), primary (n = 38), and FFPE (n = 11) samples.

We first targeted rRNA, the most abundant unwanted RNA (Fig. 4c). Adding the rRNA probe set collapsed coverage over 18S and 28S rRNA to baseline while leaving globin transcripts (HBA1, HBA2, HBB) intact. To test whether the platform could be extended to other unwanted transcripts, we added a second probe set directed against globin, a dominant transcript in blood; this depleted the globin transcripts as well. Further probes can be added to remove additional unwanted RNA, making PERD a modular depletion platform rather than a fixed rRNA kit. Depletion did not distort quantification: across 19,549 coding genes, expression with BIRT/PERD agreed with the non-depleted total-RNA library at Pearson r = 0.980 (Fig. 4d), and residual rRNA remained low across reference, primary, and FFPE samples (Fig. 4e). Integrating rRNA removal into barcoded reverse transcription therefore avoids an added cleanup while preserving both low-abundance signal and quantitative accuracy.

### Application across species and sample types

RNA sequencing is most often challenged by real-world samples in which RNA is degraded or scarce, such as archival FFPE tissue, extracellular vesicles, and long-stored blood. To test the general applicability of BIRT under these conditions, we applied it to 8,079 samples (Fig. 5a). These spanned seven species, from human, mouse, rat, and pig to fish, mosquito, and C. elegans, demonstrating that the primer chemistry transfers across organisms. Within the clinically most relevant human and mouse material, we applied BIRT across sample origins ranging from cultured cells, primary tissue, and extracted RNA to sample types that are demanding for conventional RNA-seq, including 1,813 long-stored (“aged”) PBMC samples, 1,300 extracellular-vesicle RNA samples, and FFPE tissue. Per-sample protein-coding gene detection held at a median of roughly 11,000 to 17,000 across every category, including the degraded and low-input ones (Fig. 5b). BIRT is therefore not a single-context demonstration but a method that scales across organisms and challenging sample types.

**Figure 5.**
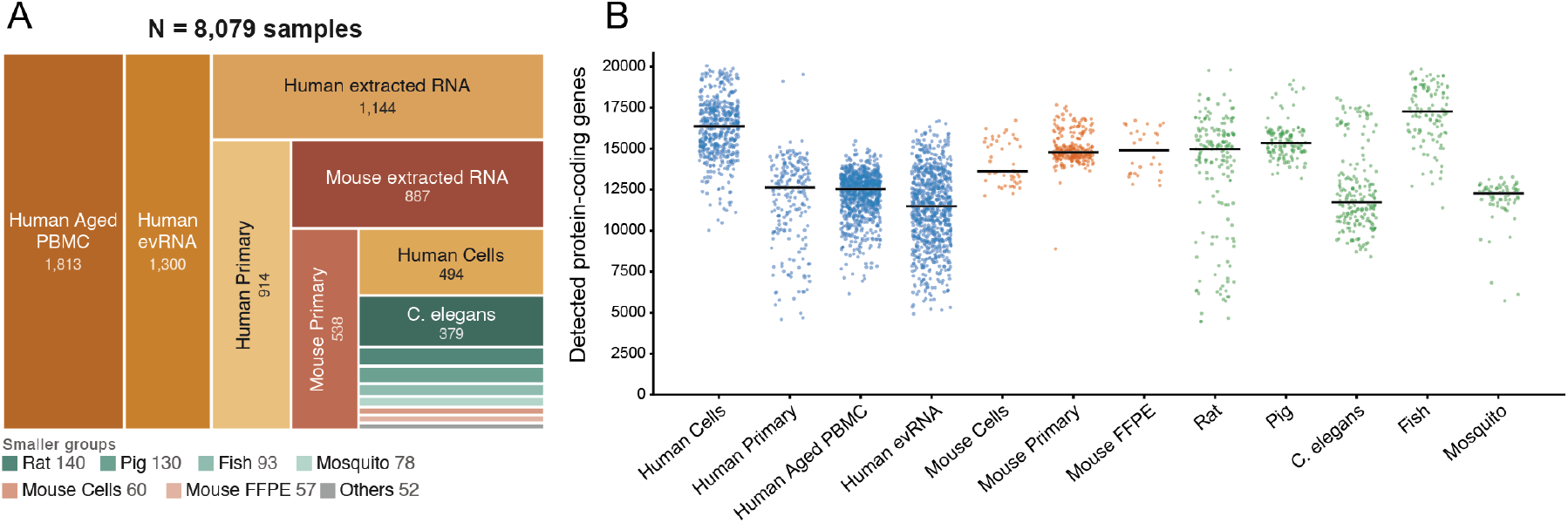
BIRT operates across species and sample types at scale. (a) Composition of 8,079 samples (counts indicated), spanning seven species and sample types including aged PBMCs, extracellular-vesicle RNA, extracted RNA, primary tissue, cultured cells, and FFPE. (b) Per-sample protein-coding gene detection by category (dot, one sample; bar, median).

## Discussion

BIRT moves the point at which sample identity, RNA provenance, and 3′-end position are recorded to the earliest step, first-strand synthesis, so that several separate problems become properties of one reaction. The barcode that enables early pooling is the same barcode that authenticates RNA-derived reads, and PERD removes unwanted RNA without adding a step. Each individual gain is modest, but they compound because they follow from a single design decision rather than from four separate fixes.

We interpret the 3′-end anchoring as a consequence of coaxial stacking, though we present this as a hypothesis rather than a directly measured mechanism. Stacking of the RNA:hexamer duplex against the primer stem is consistent with the terminal enrichment and with its dependence on a defined 3′ end, but we have not measured the thermodynamics directly, and other geometries could produce the same bias. The enrichment is also specific: BIRT marks the 3′ end of transcripts that have a discrete mature terminus, such as snoRNAs and scaRNAs, while giving full-length body coverage of mRNAs and lncRNAs; these transcripts are long and randomly fragmented, so priming falls at fragment ends set by fragmentation rather than at the transcript terminus, and terminal anchoring is not expected to be visible against the annotated end. Small nuclear RNAs reflect a reference-annotation limit rather than a biological one, their reads accumulating at a mature end that current annotations place further downstream.

The method also has bounds. BIRT does not enrich for polyadenylated RNA, so applications built on 3′-UTR isoform quantification will still prefer oligo(dT) chemistries. Within these bounds, integrating identity, authentication, priming, and depletion into reverse transcription provides a single-step route to full-length, pooled, contamination-aware total-RNA sequencing across a wide range of species and sample types.

## Methods

### Library preparation

Libraries were prepared with the NEBNext Ultra II Directional RNA workflow, replacing the free random hexamer with the BIRT primer, a hairpin oligonucleotide carrying a random hexamer at its 5′ end, a universal double-stranded stem, and a sample barcode in the loop. Because the barcode is written during first-strand synthesis, barcoded samples were pooled immediately afterward and the pool was cleaned once. For depletion, PERD blocking probes were added at the fragmentation and hybridization step; the probe panel tiles cytosolic and mitochondrial rRNA and is retargetable to other unwanted transcripts, and we additionally designed a probe set against globin. Random-hexamer (N6) control libraries were identical except that a free random hexamer replaced the BIRT primer and no barcode was added.

### Experiments

For the 3′-priming assay, synthetic RNA (120 nt) and DNA (120 and 200 nt) templates were spiked into human reference RNA (20 ng of each) and primed with the BIRT primer or a free random hexamer. For the DNA-contamination assay, human reference RNA and human male genomic DNA (Promega G147A) were mixed from 100:0 to 0:100 by mass at 100 ng total (the 90% condition being 10 ng RNA plus 90 ng DNA), with matched libraries prepared with or without TURBO DNase (Thermo Fisher AM1907); a 21-year-old primary PBMC sample was additionally processed with zero, one, or two rounds of DNase. For the pooling and low-input experiments, barcoded reference-RNA libraries were pooled after first-strand synthesis (1 to 24 barcodes), input RNA was titrated from 10 to 0.1 ng, and human/mouse mixtures were used for the barnyard species-assignment test.

### Samples

BIRT was applied to 8,079 samples spanning seven species (human, mouse, rat, pig, fish, mosquito, and C. elegans) and a range of sample types, including cultured cells, primary tissue, extracted RNA, extracellular-vesicle RNA, long-stored PBMCs, and FFPE tissue. Samples comprised human and mouse material and commercial reference RNA. All work involving human and animal samples was conducted under internal ethics approval ID 102.

### Sequencing

Libraries were sequenced paired-end (2 × 150 bp) on Illumina and Element AVITI instruments.

### Read processing

Reads were demultiplexed by BIRT sample barcode and adapter- and quality-trimmed with fastp [12] using default parameters. For contamination analysis, each read was classified as barcoded (carrying a valid BIRT barcode, RNA-derived) or non-barcoded.

### Alignment

Reads were aligned with HISAT2 [10] using default parameters: to the synthetic template sequences and the human genome for the 3′-priming assay, to the human RefSeq transcriptome (assembly GCF_000001405.40) [13] for endogenous 3′-end analysis, and to the GRCh38 human genome (GCF_000001405.40) for contamination analysis, where reads were assigned to exonic, intronic, and intergenic regions.

### 3′-end priming quantification

For each read the priming endpoint was taken as the transcript-relative position of the fragment 3′-most end. On spike-in templates, 3′-end loss was the distance from the template terminus and anchoring the fraction of priming events within a window (≤ 2 nt unless stated); the separation between BIRT and random priming was robust across 0 to 10 nt. For endogenous transcripts, distance to the annotated 3′ end was computed per read as d = transcript length − fragment 3′-most position, using first-in-pair, primary, uniquely aligned reads (samtools view -f 66 -F 2308, MAPQ ≥ 60) [11] over RefSeq non-coding (NR_) transcripts. To avoid domination by a few highly expressed genes, anchoring was computed per gene for transcripts with ≥ 20 reads and averaged across genes.

### Transcript biotype assignment

Each RefSeq transcript was assigned a biotype from the trailing descriptor of its FASTA header, with generic non-coding entries sub-classified by gene-symbol prefix (SNORD/SNORA to snoRNA, RNU to snRNA, SCARNA to scaRNA); protein-coding (NM_) transcripts were labeled mRNA.

### Contamination, pooling, and PERD analyses

DNA contamination was quantified as coverage over intronic and intergenic regions relative to exon and as the intergenic read fraction, computed separately for barcoded and non-barcoded reads. Cross-barcode reproducibility was the pairwise R^2^ of per-gene expression between barcodes; species specificity was the fraction of human/mouse reads assigned to the correct reference (barnyard). PERD depletion was assessed as coverage over 18S and 28S rRNA, and over globin, relative to retained genes, and quantitative fidelity as the Pearson correlation of per-gene expression between BIRT and the non-depleted library over 19,549 coding genes. Per-sample protein-coding gene detection (count > 0 after QC filtering) was summarized by sample category as the median.

### Statistics

Correlations are Pearson. In dot plots, each dot is one sample and the bar is the median; group sizes (N) are given in the figure legends.

## Data and code availability

Sequencing data and analysis code are available from the corresponding authors upon reasonable request.

